# A computational method for the investigation of multistable systems and its application to genetic switches

**DOI:** 10.1101/088005

**Authors:** M. Leon, M. L. Woods, A.J.H. Fedorec, C.P. Barnes

## Abstract

Genetic switches exhibit multistability, form the basis of epigenetic memory, and are found in natural decision making systems, such as cell fate determination in developmental pathways. Synthetic genetic switches can be used for recording the presence of different environmental signals, for changing phenotype using synthetic inputs and as building blocks for higher-level sequential logic circuits. Understanding how multistable switches can be constructed and how they function within larger biological systems is therefore key to synthetic biology. Here we present a new computational tool, called StabilityFinder, that takes advantage of sequential Monte Carlo methods to identify regions of parameter space capable of producing multistable behaviour, while handling uncertainty in biochemical rate constants and initial conditions. The algorithm works by clustering trajectories in phase space, and iteratively minimizing a distance metric. Here we examine a collection of models of genetic switches, ranging from the deterministic Gardner toggle switch to stochastic models containing different positive feedback connections. We uncover the design principles behind making bistable, tristable and quadristable switches, and find that rate of gene expression is a key parameter. We demonstrate the ability of the framework to examine more complex systems and examine the design principles of a three gene switch. Our framework allows us to relax the assumptions that are often used in genetic switch models and we show that more complex abstractions are still capable of multistable behaviour. Our results suggest many ways in which genetic switches can be enhanced and offer designs for the construction of novel switches. Our analysis also highlights subtle changes in correlation of experimentally tunable parameters that can lead to bifurcations in deterministic and stochastic systems. Overall we demonstrate that StabilityFinder will be a valuable tool in the future design and construction of novel gene networks.

## 1 Background

Synthetic biology has seen the development of many simple gene circuits such as switches [1, 2, 3, 4, 5, 6], oscillators [7, 8, 9] and pulse generators [10]. Larger systems have been constructed [11], but the leap from building low-level circuits to assembling them into complex networks is still a major challenge [12, 13]. Efforts to do so are plagued by circuit crosstalk, retroactivity, chassis loading effects, and cellular noise, which can render synthetic networks non-functional *in vivo* [14, 15]. Although standardization and better part design can partially lower this barrier [12, 16, 17, 18, 19], design processes that enable the informed selection of appropriate parts are crucial [20, 21, 11].

One of the foundational constructs in synthetic biology is the genetic toggle switch. The toggle switch consists of a set of transcription factors that mutually repress each other [1, 22, 23, 24]. Genetic switches play a major role in binary cell fate decisions such as stem cell differentiation, as they are capable of exhibiting bistable behaviour, which gives rise to the existence of two distinct phenotypic states. This allows populations of cells to maintain a heterogeneous response to environmental cues and can increase fitness by bet-hedging [25]. Switches are powerful building blocks; they underlie electronics and logic systems, and have great potential in synthetic biology. The genetic toggle switch has been used for a number of applications including the construction of a synthetic genetic clock [22], the regulation of mammalian gene expression [5, 2], the development of a predictable genetic timer [26], and the formation of biofilms in response to engineered stimuli [27].

The stability of the toggle switch has been investigated extensively in the literature, but the conclusions drawn vary according to model abstraction. Numerous studies have concluded that cooperativity is a necessary condition for bistability to arise [1, 28, 29, 30, 31]. However, Lipshtat *et al*. found that stochastic effects can give rise to bistability even without cooperativity [32]. In another study, Ma *et al*. found that stochastic fluctuations can stabilize the unstable steady state in the deterministic system, giving rise to tristability [33]. In addition, Biancalani *et al*. identified multiplicative noise as the source of bistability in the stochastic case [34]. As is clear from the above, there is yet to exist a consensus on the stability a switch is capable of, and the most appropriate method of modelling it. Most of these studies assumed the quasi-steady state approximation (QSSA) [35], which cannot always be assumed to hold *in vivo* [36].

In terms of system design, extensions of the basic toggle switch motif, including additional positive feedback mechanisms, have been investigated [37, 38], and optimization methods have been used to identify topologies and parameter values for bistable and tristable genetic switches [39, 40, 41, 42]. For stochastic switch design, control theoretic approaches [43], and simulation-based frameworks [44], have been developed. However, none of these existing approaches can be be applied to reasonably sized models, under the assumption of deterministic and stochastic dynamics, and identify regions of parameter space for which switching occurs, which we argue is critical in designing systems under considerable uncertainty.

Here we present a computational framework based on sequential Monte Carlo [45] that can determine the parameter region for a given model to produce a given number of (stable) steady states. Uniquely, multistable parameter regions can be identified for both deterministic and stochastic systems, and also complex models with many parameters, thus removing the need for simplifying assumptions. Our framework can be used for comparing the conclusions drawn by various modelling approaches and thus provides a way to investigate appropriate abstractions. This framework is available as a Python package, called StabilityFinder. We investigate genetic toggle switches and uncover the design principles behind making bistable and tristable switches (all models used in this study are summarised in Table 1.) We find that both production and degradation rates of transcription factors are key parameters for bistability, and outline how the addition of positive autoregulation, combined with particular parameter combinations, can create multistable switching behaviour. Finally we demonstrate the ability of the framework to examine more complex systems and examine the design principles of a three gene switch. These examples demonstrate that StabilityFinder will be a valuable tool in the future design and construction of novel gene networks.

**Table 1:** Summary of the models used in this study.

|  | Model | Autoregulation (+ve) | Steady states | Reference |
| --- | --- | --- | --- | --- |
| Deterministic | Gardner | None | 2 | [1] |
|  | LU-CS | None | 2 | [38] |
|  | LU-SP | Single | 2 | this study |
|  | LU-DP | Double | 2, 3, 4 | [38] |
|  | Three gene switch | Triple | 6 | this study |
| Stochastic | Gardner | None | 2 | [1] |
|  | MA-CS | None | 2, 3 | this study |
|  | MA-DP | Double | 2, 3 | this study |

## 2 Methods

StabilityFinder is based on a statistical inference method which combines approximate Bayesian computation (ABC) with sequential Monte Carlo [46]. This simulation-based method uses an iterative process to arrive at a distribution of parameter values that can give rise to observed data or a desired system behaviour [44]. ABC methods are used for inferring the posterior distribution in cases where the likelihood is intractable or is too computationally expensive to evaluate. Instead of computing the likelihood, ABC methods simulate the data and then compare the simulated and observed data through a distance function [46]. Given the prior distribution *π*(*θ*) we can approximate the posterior distribution, *π*(*θ* | *x*) ∝ *f*(*x* | *θ*)*π*(*θ*), where *f*(*x* | *θ*) is the likelihood of a parameter, *θ*, given the data, x. There are a number of different variations of the ABC algorithm depending on how the approximate posterior distribution is sampled.

The simplest ABC algorithm is the ABC rejection sampler [47]. In this method, parameters are sampled from the prior and data simulated through the data generating model. For each simulated data set, *x*\*, a distance from that of the desired behaviour is calculated, *ρ*(*x*, y*), and if greater than a threshold, ∈, the sample is rejected, otherwise it is accepted. The main disadvantage of this method is that if the prior distribution is very different from the posterior, the acceptance rate is very low [46]. An alternative method is the ABC Markov Chain Monte Carlo (MCMC) [48]. The disadvantage of this method is that if it gets stuck in an area of low probability it can be very slow to converge [49]. The method used here is based on sequential Monte Carlo, which avoids both issues faced by the rejection and MCMC methods. It propagates the prior through a series of intermediate distributions in order to arrive at an approximation of the posterior. The tolerance, ∈, for the distance of the simulated data to the desired data is made smaller at each iteration. When ∈ is sufficiently small, the result will approximate the posterior distribution [46].

**Algorithm 1.**
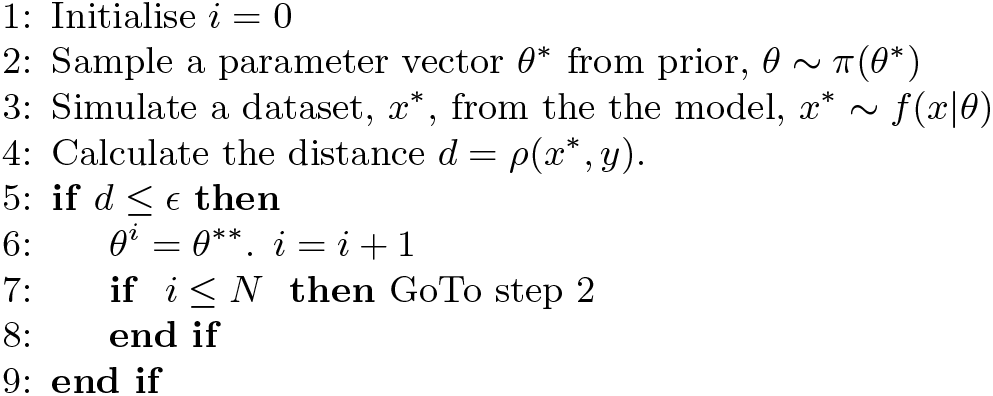
ABC rejection algorithm to generate samples {*θ^i^*; *i* ≤ *N*} from *π*(*θ*|*ρ*(*x, y*) ≤ ∈)

To investigate the multistable behaviour of systems, a number of extensions to existing approaches are required. For a given set of parameter values, sample points are taken across initial conditions using latin hypercube sampling [50], and the ensemble system simulated in time until steady state. The distance function in ABC is replaced by a distance on the desired stability of the simulated model. To do this we cluster the steady state coordinates using K-means clustering [51] and use the Gap statistic to determine the number of clusters [52]. At each iteration, the number of steady states is determined by the number of clusters in phase space. A particle is accepted only if the number of clusters present is within an acceptable distance from the threshold e. The algorithm is summarised below.

This algorithm is available as a Python package, called StabilityFinder. The user provides an SBML model file [53, 54] and an input file that contains all the necessary information to run the algorithm, including the desired stability and the final tolerance, ∈, for the distance from the desired behaviour necessary for the algorithm to terminate. The flow of execution is illustrated in Figure 1. Since the algorithm is computationally intensive, all deterministic and stochastic simulations are performed using algorithms implemented on Graphics Processing Units (GPUs), which are used for mutli-threaded computation [55]. The algorithm returns the final accepted particles and their associated weights, as well as the initial conditions sampled and the steady state values obtained. The final accepted particles can be used to study the characteristics of the posterior distribution. The sampled initial conditions and the resulting steady state values can be used to study the basins of attraction of the system.

**Figure 1.**
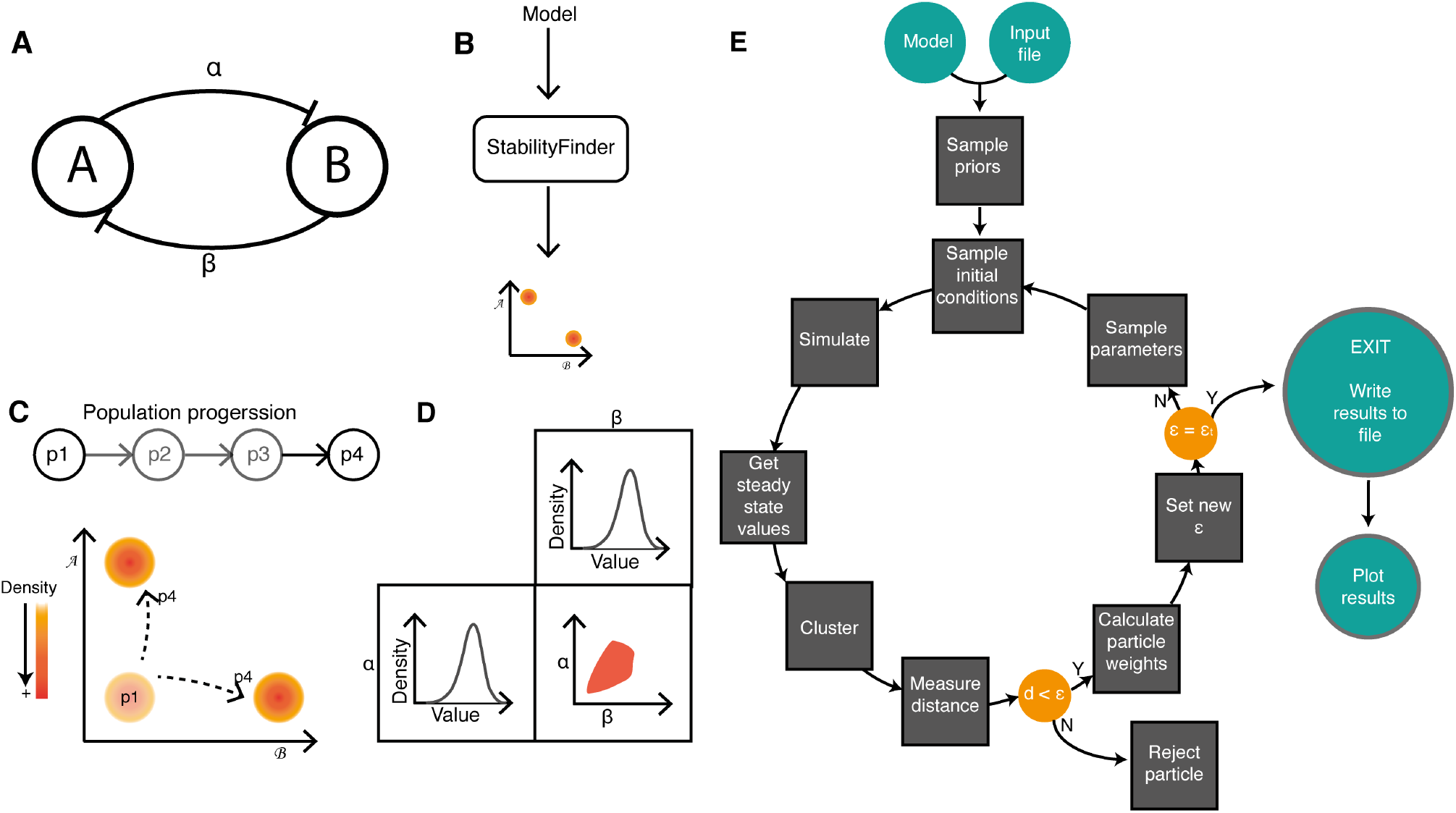
Using sequential Monte Carlo to examine system stability. (A,B) The algorithm takes as input a model and a desired stability structure in phase space. (C) The system is evolved to achieve the stability of choice via intermediate populations. In this example model, there are two species and two parameters. For the model to be considered bistable, the phase plot of the two species of interest must have two distinct clusters, as shown in (C). (D) The parameter posterior distribution for the model to achieve the desired behaviour are given as an output. (E) Flow chart of the Python package, StabilityFinder, which implements the algorithm.

## 3 Results and Discussion

### 3.1 The Gardner switch under deterministic and stochastic dynamics

The first synthetic genetic toggle switch was constructed in *E. coli* by Gardner *et al*. and consisted of two mutually repressing transcription factors [1]. The model used to design and interpret the system is shown in Figure 2A, and in the deterministic case is defined by the following ODEs

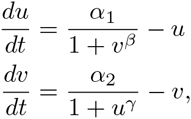

where *u* is the concentration of repressor 1, *v* the concentration of repressor 2, *α*_1_ and *α*_2_ denote the effective rates of synthesis of repressors 1 and 2 respectively, and *β* and *γ* are the cooperativity of repression of promoter 1 and of repressor 2 respectively. Gardner *et al* studied the deterministic case and concluded that there are two conditions for bistability for this model; that *a\* and *a_2_* are balanced and that *β*, *γ* >1 [1]. In order to test StabilityFinder we used it to find the posterior distribution for which this model exhibits bistable behaviour. We therefore set the desired behaviour to two (stable) steady states, and using a wide range of values as priors as shown in the Supplementary Information, we used StabilityFinder to find the parameter values necessary for bistability to occur. The posterior distribution calculated by StabilityFinder for the Gardner deterministic case is shown in Figure 2C. These results agree with the results reported by Gardner *et al*. [1]. For this switch to be bistable *α*_1_ and *α*_2_ must be balanced while *β* and *γ* must both be > 1, as can be seen in the marginal distributions of *β* and *γ* in Figure 2C.

**Figure 2.**
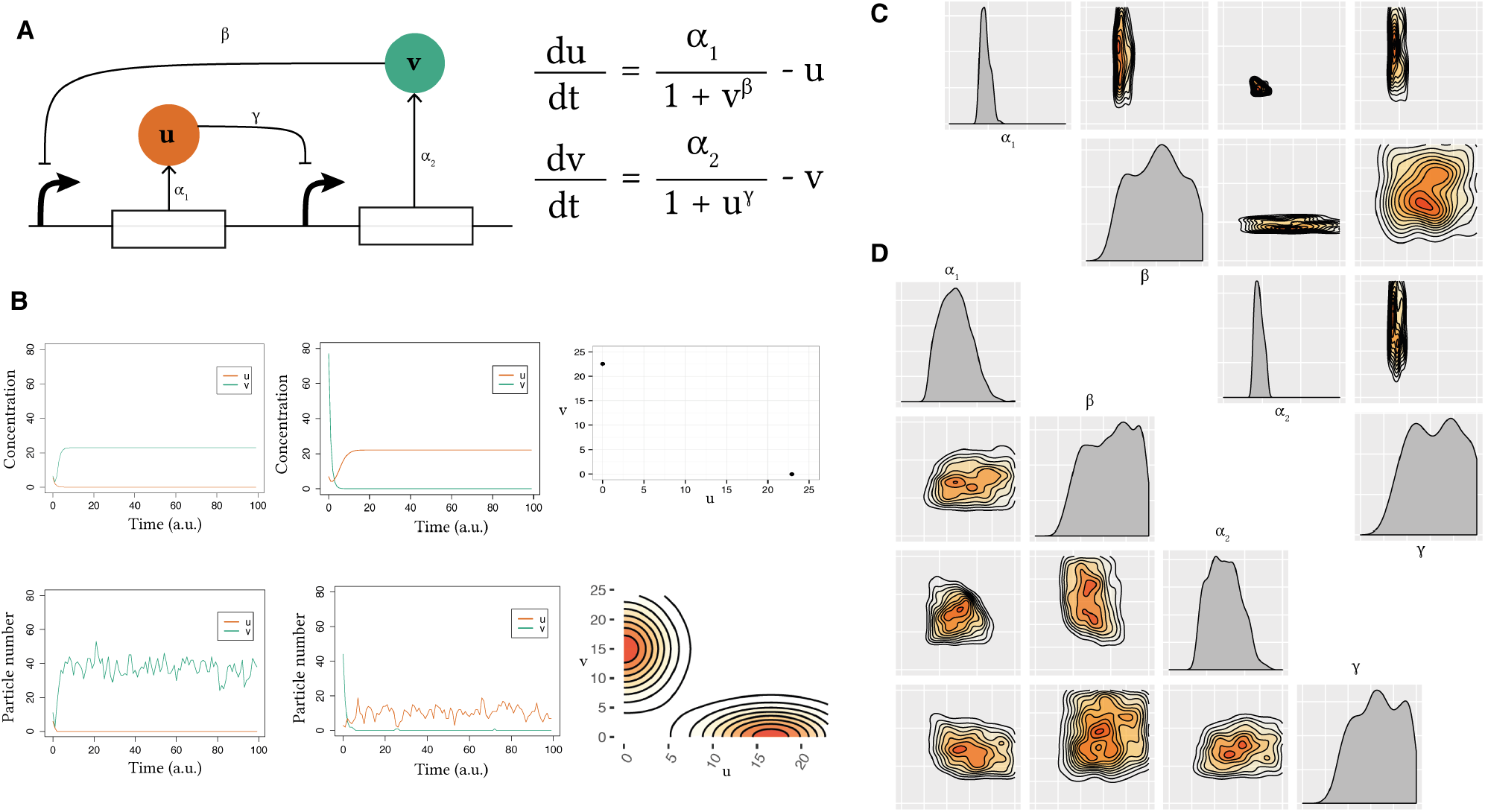
Analysis of the Gardner switch model. (A) The model consists of two mutually repressing transcription factors. It can be reduced to a two-equation system, where *u* and *v* are the concentrations of transcription factors 1 and 2, *α*_1_, *α*_2_ are their effective rates of synthesis, and *β*, *γ* represent the cooperativity of each promoter. (B) Two samples of deterministic simulated time courses of the Gardner switch and the resulting phase plot and two samples of time courses of the stochastic simulations and the resulting stationary distributions. (C,D) The deterministic (C) and stochastic (D) posterior distributions. These include the one dimensional marginal density plots on the diagonal, and the two dimensional marginal distributions on the off-diagonal. All densities are plotted on the prior range.

**Algorithm 2.**
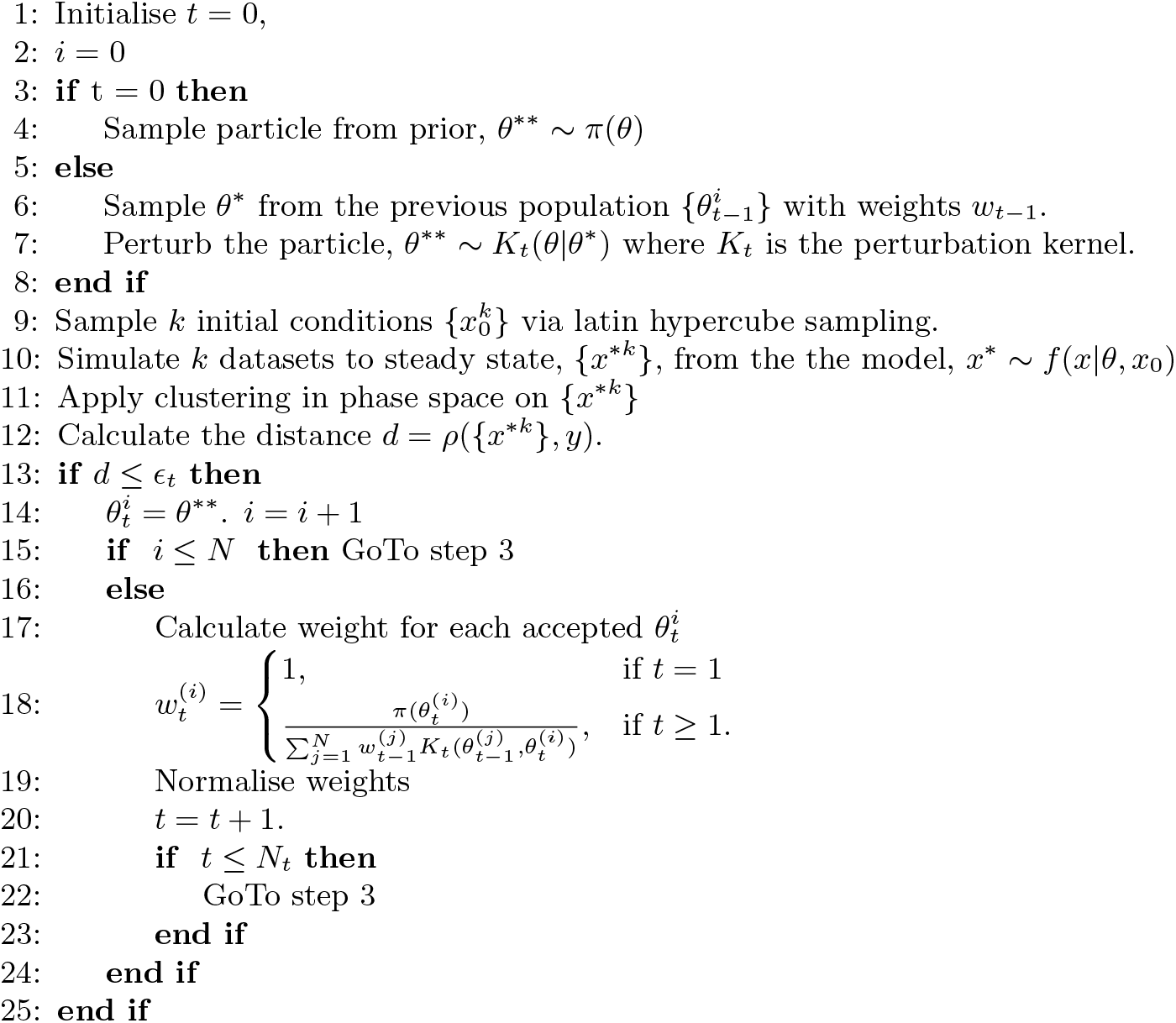
Stability Finder algorithm

We next applied StabilityFinder to the case of the Gardner switch under stochastic dynamics using the same priors as the deterministic case, and again searched the parameter space for bistable behaviour. The posterior is shown in Figure 2D. We can see that the conditions on the parameters required for bistability in the deterministic case generally still stand in the stochastic case. There appears to be slightly looser requirements on the parameters of the stochastic model (wider marginal distributions), which is expected due to the nature of clustering deterministic steady states versus stochastic steady states. The Gap statistic is used in the case of the stochastic case, as it is capable of dealing with noisier data whereas a simpler and faster algorithm is used for clustering the deterministic solutions (see Supplementary Information). These results demonstrate that StabilityFinder can be used to find the parameter values that can produce a desired stability and allow us to confidently apply the methodology to more complex models.

### 3.2 Repressor degradation rates are key for achieving bistablity

We next analyzed an extension of the Gardner switch model previously studied by Lu *et. al*. [38]. They considered two types of switches, the classic switch consisting of two mutually repressing transcription factors (model LU-CS), as well as a switch with double positive autoregulation (model LU-DP). The LU-CS switch was found to be bistable given the set of parameters used, while the LU-DP switch was found to be tristable [38]. The classical model used in their study is given by the following system of ODEs

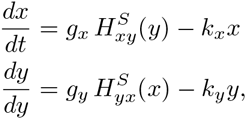

where

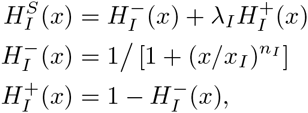

and *g_I_* represents the production rate, *k_I_* the degradation rate, *n_I_* the Hill coefficient, *x_I_* the Hill threshold concentration and *λ_I_* the fold change of the transcription rates, and *I* ∈ {*xy, yx, xx, yy*}. (see Supplementary Information for the details of all models used).

For the parameter values used in the Lu study, the classical switch exhibits three steady states (Figure 3), two of which are stable and one is unstable. Using StabilityFinder with priors centred around the parameter values used in the original paper (see Supplementary Information), we can identify the most important parameters for achieving bistability. The posterior distribution of these models are shown in Figure 3A. We find that the parameters representing the rates of degradation of the transcription factors in the system (*k_x_, k_y_*) must both be large in relation to the prior range, and approximately equal, for bistability to occur. Protein degradation rates have been shown to be important for many system behaviours including oscillations [7, 56, 57]. We also find that the steady states of the LU-CS model are symmetric: the values for the dominant and repressed species are equivalent in both steady states.

**Figure 3.**
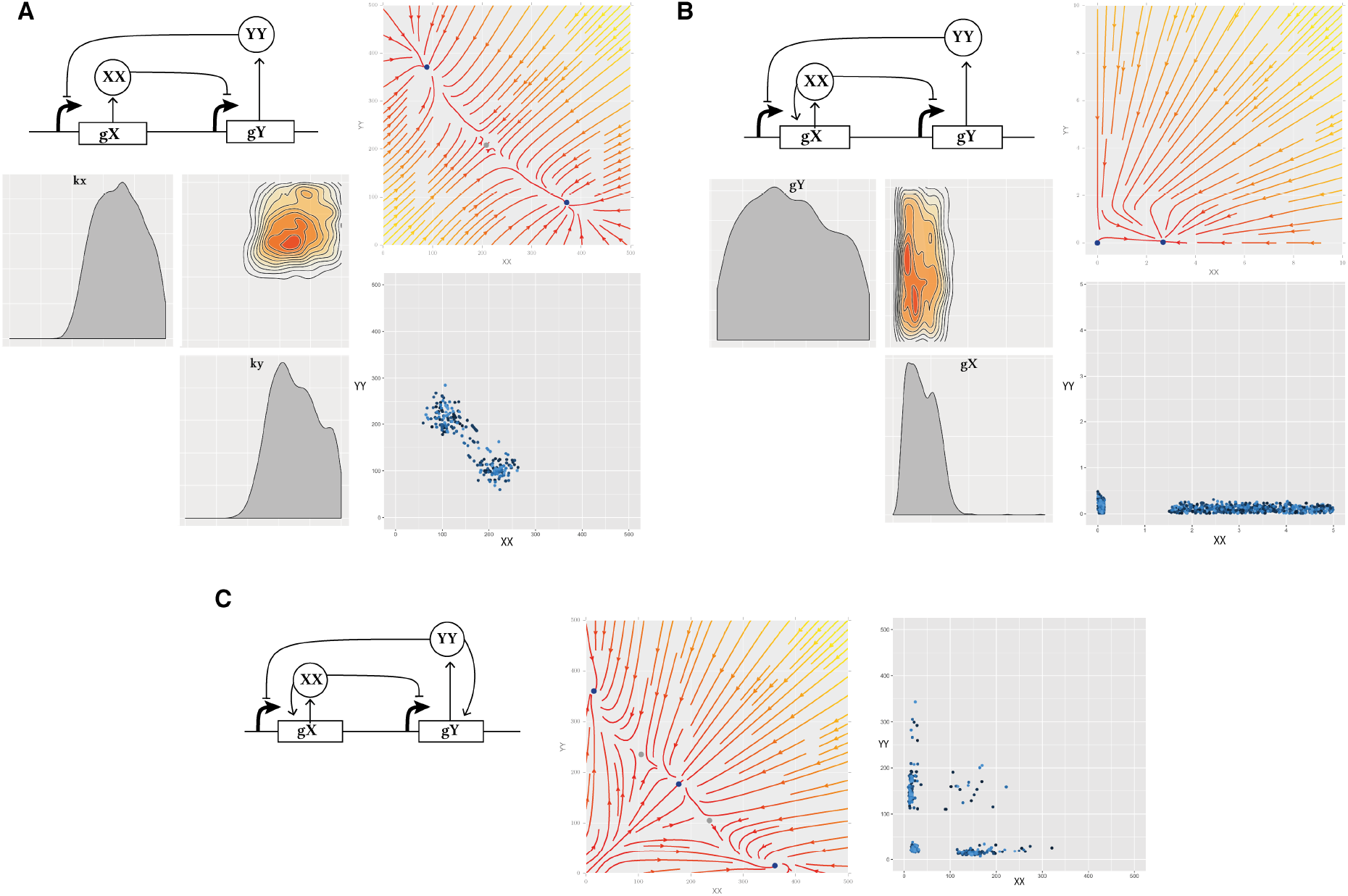
The three variants of the deterministic Lu models. (A) The classic switch with no autoregulation is bistable as shown in the stream plot and the phase plot. In the stream plot, the colours indicate the magnitude of the vectors, with yellow indicating high and red low values. The blue points represent stable steady states and the grey points represent unstable steady states. The phase plot shows the steady state values for 100 particles at the final population. Each particle is represented by a different shade of blue. The most restricted parameters for this behaviour are the degradation rates (*k_x_* and *k_y_*), which both have to be high while the net protein production for *X* and *Y* must be balanced. (B) The extended Lu model with a single positive autoregulation on *X*. This model is bistable when the production rate of *X*, *g_x_*, is small. (C) The Lu model with double positive autoregulation is tristable as shown in the stream plot and the phase plot. We find two types of tristable behaviour, one where the third steady state is zero-zero and one where the third state is high.

### 3.3 The addition of symmetric and asymmetric positive autoregulation

It is known that the addition of positive autoregulation to the classical toggle switch can induce tristability [58, 38, 37]. Here we investigate the interplay of positive autoregulation on the values of the other parameters in the model. We extended the analysis presented in Lu *et al*. by including the switch with single positive autoregulation (model LU-SP), where a single positive autoregulatory feedback is present on one of the genes. This system topology has also been constructed previously [23, 59]. The advantage of using StabilityFinder over traditional bifurcation analysis is that the full parameter space is explored rather than solving the system for a single set of parameters. This allows us to deduce model properties that could not otherwise be identified. Robustness to parameter fluctuations can be explored, as well as parameter correlations and restrictions on the values they can take while still producing the desired behaviour.

The LU-DP model is given by

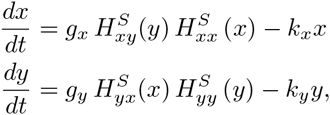

whereas the LU-SP switch is modelled using the following ODE system

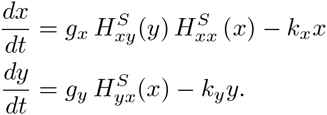

We find that the switch with single positive autoregulation is capable of bistable behaviour as seen in Figure 3B, but this is only possible when the strength of the promoter under positive autoregulation, *g_x_*, is small. There appear to be no such constraints on the strength of the original, unmodified, promoter, *g_y_*. We also find that unlike the LU-CS and LU-DP models, the steady states of the bistable LU-SP are not symmetric. The levels of Y are around zero and always lower than the levels of X. The levels of both are lower than those found in the LU-CS and LU-DP models.

Upon examination of the LU-DP model, we also find that tristability in the switch is relatively robust, as this phenotype is found across a large range of parameter values, with no parameters strongly constrained (see Supplementary Information), but the two parameters for gene expression, *g_x_* and *g_y_* tend to be small compared to the priors. Two types of tristable behaviour are identified, one where the third steady state is at (0,0) and and one where the third steady state has non-zero values. This result agrees with previous work where it was found that a switch can exhibit two kinds of tristability, one in which the third steady state is high (III*_H_*) and one in which it is low (III*_L_*) [37].

### 3.4 Design principles for a switch capable of two, three and four steady states.

The LU-DP switch is capable of both bistable and tristable behaviour as well as four coexisting states under deterministic dynamics (quadristability) [37]. It is of great interest to understand the conditions under which these three behaviours occur. We carried out a bifurcation analysis of the DP switch using the PyDSTool [60] in order to get an indication of the stabilities this model is capable of, and at which parameter ranges these are found (Figure 4B). This shows that by varying the parameter for gene expression (*g_x_*) while all other parameters remain constant we can obtain all three behaviours. We find that if 100 ≤ *g_x_* ≤ 120 the system exhibits four steady states, if 9 ≤ *g_x_* ≤ 10 the system is tristable and if 10 ≤ *g_x_* ≤ 100 the system is bistable (see Supplementary Information).

**Figure 4.**
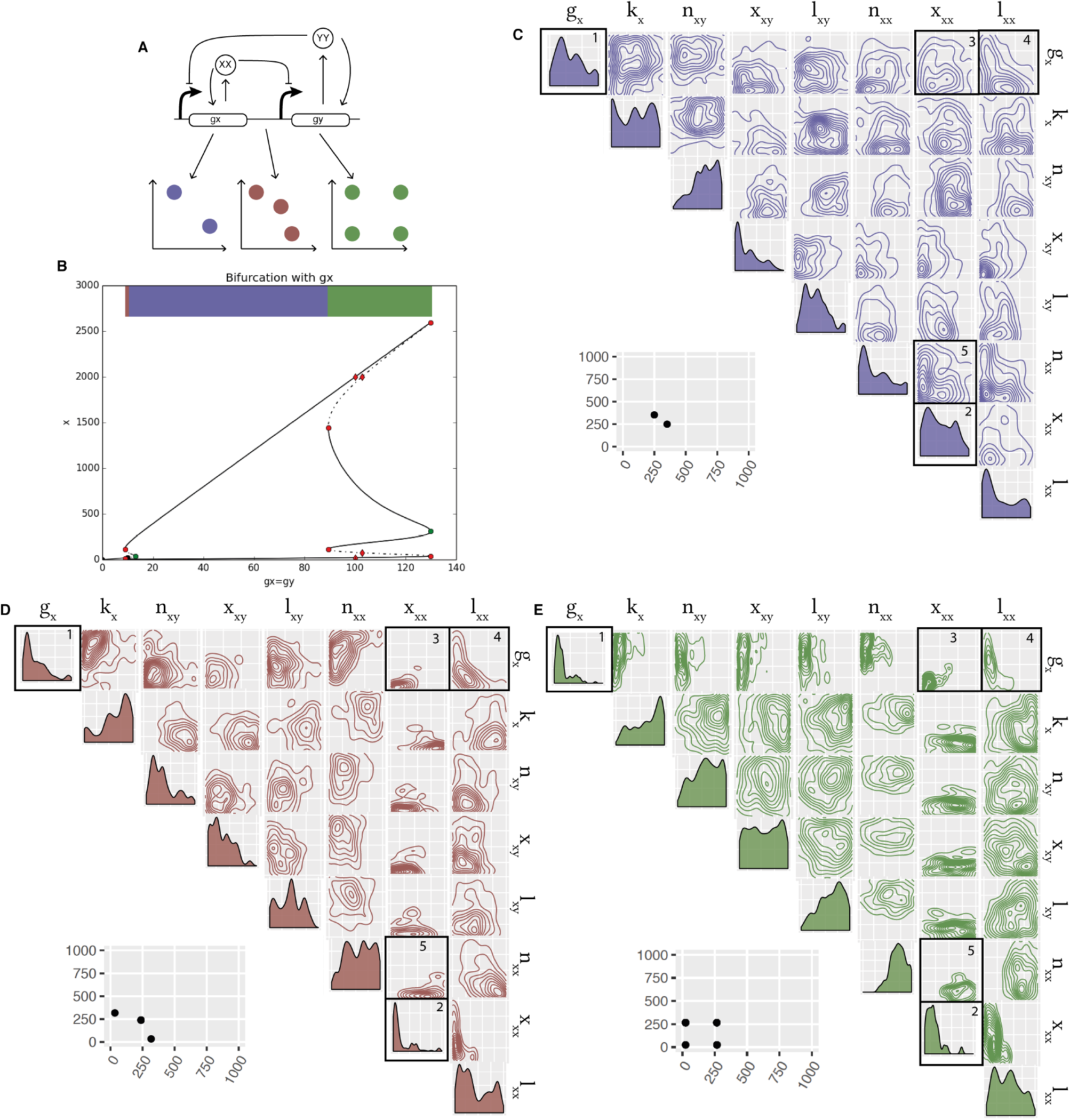
Design principles of multistable switches. (A) Using the Lu model with added positive autoregulation we uncover the design principles dictating if a switch will be bistable, tristable, or quadristable. (B) An initial bifurcation analysis of the LU-DP switch uncovers the stabilities it is capable when varying the parameter for gene expression (while keeping all other parameters fixed). (C-E) By considering the bivariate distributions of the parameters we can uncover the differences in the parameters of a bistable switch compared to a tristable switch, compared to a quadristable switch. The posterior distribution of the bistable switch is shown in purple, the tristable switch in red and the quadristable switch in green, all plotted on the prior ranges. The bivariate distributions for which a difference is observed between the stabilities are in black boxes. An example of the phenotype (phase plot) from each switch is shown next to the corresponding posterior distribution. Parameter legend key: *g_x_, g_y_* production rates; *k_x_, k_y_* degradation rates; *n_xy_, n_yx_* Hill coefficients; *x_xy_, x_yx_* Hill threshold concentrations; *l_xy_, l_yx_* transcription rate fold change; *n_xx_, n_yy_* autoregulation Hill coefficients; *x_xx_, x_yy_* autoregulation Hill threshold concentrations; *l_xx_, l_yy_* autoregulation transcription rate fold change.

Using StabilityFinder we obtained posterior distributions for the bistable, tristable and quadristable phenotypes (Figure 4). Upon examination of the these distributions, we observe that a subset of the parameter values are different for the three behaviours (Figure 4C), although the differences are surprisingly subtle. We find differences in the univariate distribution of the parameters for gene expression, *g_x_*, as highlighted in Figure 4C, box 1. This parameter must be small for four steady states to occur but there are no such restraints for a bistable or a tristable switch. Furthermore, parameter *x_xx_* (the dissociation constant for autoregulation) must be small for tristable and quadristable behaviour to be achieved, but there are no such restraints for a bistable switch, as can be seen in Figure 4C, box 2.

Additionally, we find a difference in the bivariate distributions in the posterior. Most notably, we find that parameters *x_xx_* and *g_x_* are tightly constrained in the tristable and quadristable cases, where both parameters are required to be small, but less so in the bistable case (Figure 4C, box 3). Another notable difference is between parameters *x_xx_* and *n_xx_* shown in Figure 4C, box 5, where they are constrained in the tristable and quadristable cases but not the bistable case. Interestingly, we also find parameter correlations conserved between the three behaviours, as seen in Figure 4C, box 4, where parameters *l_xx_* and *g_x_*, (positive autoregulation and gene expression) are negatively correlated in both cases. This highlights the importance of treating unknown parameters as distributions rather than fixed values when studying complex system behaviour. These ensemble based methods are capable of uncovering not only the ranges and values required for certain behaviour, but also the correlations between parameters, which would be missed by optimisation methods.

### 3.5 Bistability and tristability using more realistic abstractions

In order to study the switch system in the most realistic way, we avoid using the quasi-steady state approximation (QSSA) that is often used in modelling the toggle switch. Under the assumption of mass action kinetics, the two-equation system becomes a system of 14 equations and 9 parameters in the classical switch case. These are shown in the Supplementary Information. In the cellular environment stochastic effects can be non-negligible and should therefore be taken into account when trying to elucidate the behaviours that a system is capable of. Therefore, we model these switches using stochastic dynamics.

We find that under stochastic dynamics, both the simple switch, CS-MA, and positive autoregulation switch, DP-MA, are capable of bistable and tristable behaviour (Figure 5). The fact that tristability can occur in the classical model is consistent with the effect of small molecule numbers; if gene expression remains low, it provides the opportunity for small number effects to be observed, and the third unstable steady state to stabilise [33]. To verify the robustness of the tristability found in the stochastic case, we re-sampled the posterior distributions, simulated to steady state and confirmed tristable behaviour. As can be seen in Figure 5, differences in the parameter values are observed between the bistable and tristable switches, in both CS-MA and DP-MA models. The design principles for both the CS-MA model and the DP-MA model are summarised in Table 2.

**Figure 5.**
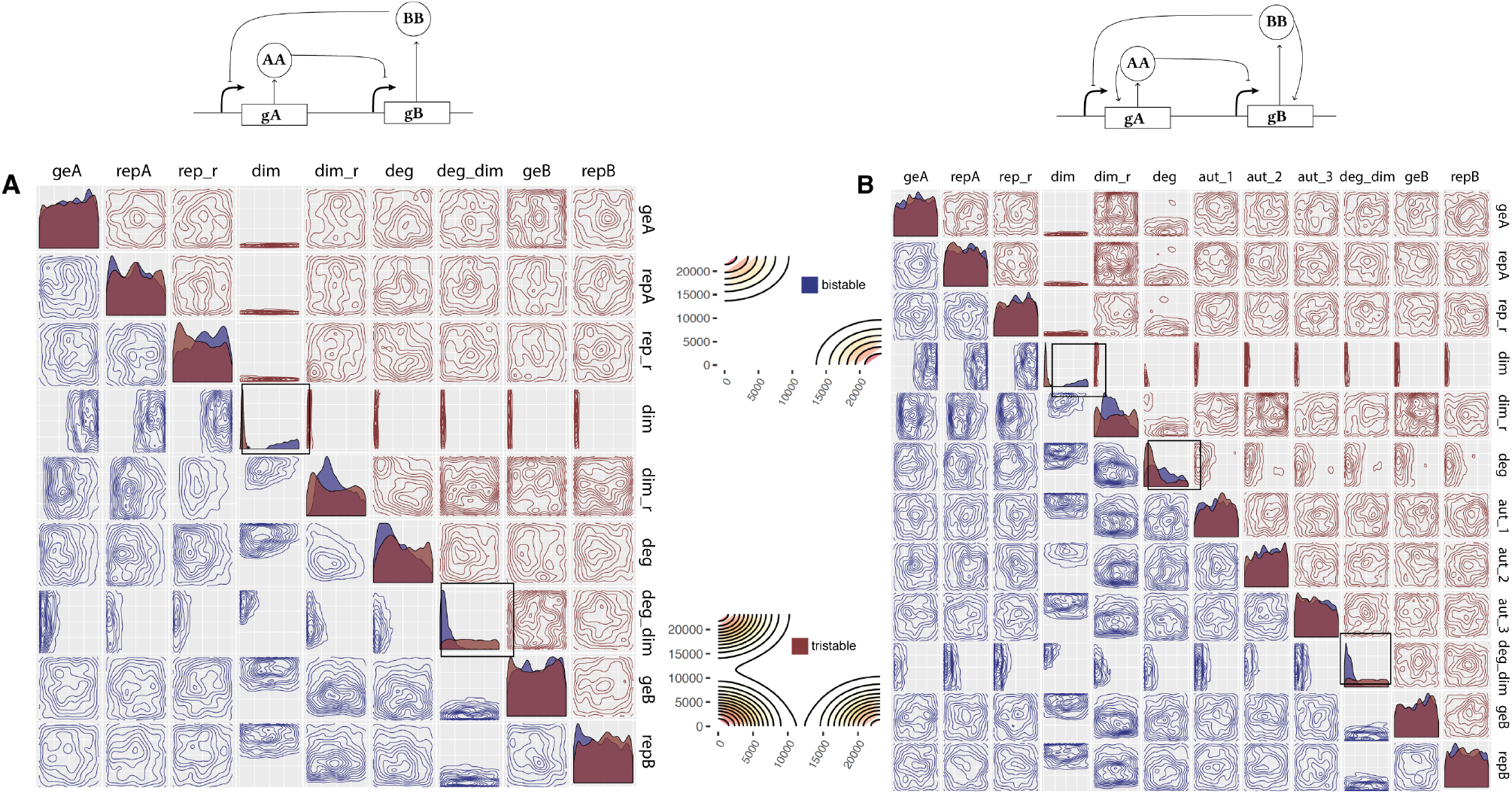
Tristability is possible in the mass action toggle switch models only when simulated stochastically. (A) The simple toggle switch with no autoregulation can be both bistable and tristable. The two posteriors are shown, plotted on the prior ranges, where the posterior distribution of the bistable switch is shown in blue and of the tristable switch in red. From the posterior distribution we can deduce the the dimerization parameter must be small for tristability to occur but large for bistability. (B) The switch with double positive autoregulation and its posterior distributions for the bistable and tristable case.

**Table 2:** Design principles of bistable and tristable switches. Differences are observed between the parameter values of the bistable and tristable CS-MA and DP-MA switches.

|  | CS-MA |  | DP-MA |  |
| --- | --- | --- | --- | --- |
|  | *Bistable* | *Tristable* | *Bistable* | *Tristable* |
| dimerisation | High | Low | High | Low |
| protein degradation | - | - | - | Low |
| dimer degradation | Low | - | Low | - |

### 3.6 Achieving higher multistability

To further demonstrate the flexibility of our framework we investigated a system capable of higher stabilities. Multistability is found in some differentiating pathways, such as the myeloid differentiation pathway [61, 62]. We allow for these more complex dynamics by extending the LU-DP model by adding another gene, making it a three gene switch. This new system is depicted in Figure 6A. In StabilityFinder we look for six steady states, the output being in nodes *X* and *Y* and use the priors shown in the Supplementary information. We successfully find that the system is capable of six steady states, as shown in Figure 6C. Consistent with the LU-DP switch capable of 2, 3 and 4 steady states, we find that the steady states are symmetric (Figure 6C). Each of the six steady states exists in symmetry with another one, in tightly constrained regions. We find that the most constrained parameters for this behaviour are again the degradation rate of the proteins, *k_x_*. If they are too large or too small the system will not exhibit hexastability. Additionally we find that the Hill coefficients for the repressors, *n_xy_*, are constrained to be smaller than 1.5 as seen in Figure 6D. This example demonstrates that Stability Finder can be used to elucidate the dynamics of more complex network architectures, which will be key to the successful design and construction of novel gene networks as synthetic biology advances.

**Figure 6.**
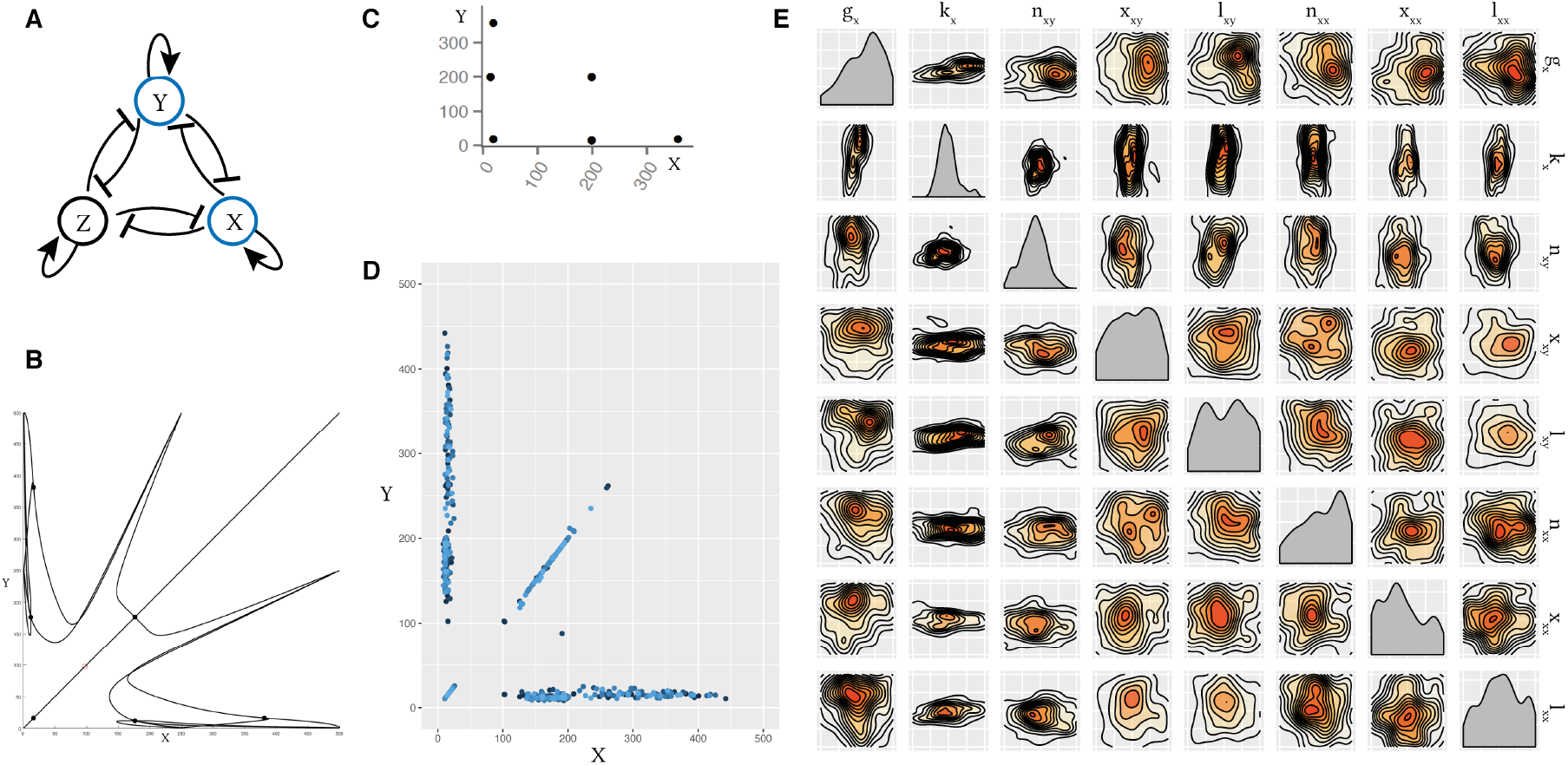
The three-node mutual repression model, with added positive auto-regulation on each node. (A) The model is studied in two dimensions using StabilityFinder, with clustering performed on the levels of *X* and *Y*. (B) Null clines and steady states of the three-node switch. (C) An example phase plot from the posterior of the three-node switch and (D) the phase plot of 100 particles from the final model. Each particle is represented by a different shade of blue. (E) The posterior distribution of the 6-steady state three-node system plotted on the prior ranges. Parameters for degradation, *k_x_*, and the Hill coefficient, *n_xy_*, are the most constrained. Parameter legend key: *g_x_* production rate; *k_x_* degradation rate; *n_xy_* Hill coefficient; *x_xy_* Hill threshold concentration; *l_xy_* transcription rate fold change; *n_xx_* autoregulation Hill coefficient; *x_xx_* autoregulation Hill threshold concentration; *l_xx_* autoregulation transcription rate fold change.

## 4 Conclusions

We have developed an algorithm that can identify the parameter regions necessary for a model to achieve a given number of stable steady states. The novelty in our framework over existing methodology is that complex models can be analyzed assuming both deterministic and stochastic dynamics. We have shown that the algorithm can be used to infer the parameter ranges that give rise to specific behaviour in various models. We uncovered the design principles that make a bistable, a tristable and a quadristable switch. We also found that a three-node switch is capable of hexastability. Importantly, we removed assumptions made to simplify the switch models and showed that they are still capable of bistable and tristable behaviour.

Although we only examined models containing combined transcription and translation, our approach could be applied to any models of switching behaviour, including more detailed kinetic models and more complex multistable switches that exist in natural biological systems, such as developmental pathways. We also limited our framework to the objective behaviour of a given number of stable steady states. However, this approach is extremely flexible, and could be extended to find systems with a given switching rate, or systems robust to a particular set of perturbations, both of which could be of great importance for building more complex genetic circuits.

One limitation of our approach is that we cannot rule out a specific behaviour; it is always possible that some part of parameter space remains unexplored, or because the search space must be predefined, interesting regions are not included in the search. In the Bayesian sense, this predefined space is the prior distribution for the parameters that give rise to the stability under investigation. In principle, once our knowledge of these biochemical rate constants grows, we can incorporate these data into the prior regions for exploration. Another limitation is that of scalability. Our framework can currently be applied to small and medium size gene networks since the computational time is exponential in size, whereas optimization methods are more scalable [39, 40, 41]. This is a manifestation of a general tradeoff between finding an optimal value and exploring a parameter space. However, we argue that for current and relevant problems in synthetic biology, this computational burden is acceptable.

Approaches based on parameter space exploration are indispensible tools for providing understanding of general system properties and guiding more detailed experimental and theoretical studies. They will also be key for the design and construction of synthetic gene networks. By selecting standardized parts accordingly, such as promoters, RBS sequences and other untranslated regions [63, 64, 18, 65], *in vivo* systems can be matched to parameter regions with a high probability of function.

More generally our results highlight that changing the level of abstraction, in addition to the modification of the feedback structure and parameter values, can significantly alter the qualitative behaviour of a system model. These results advocate the need for a programme of experimental work, combined with systems modelling, to understand the rules of thumb for abstraction in model based design of synthetic biological systems.

## Acknowledgements

All the authors would like to acknowledge that the work presented here made use of the Emerald High Performance Computing facility made available by the Centre for Innovation. The Centre is formed by the universities of Oxford, Southampton, Bristol, and University College London in partnership with the STFC Rutherford-Appleton Laboratory.

